# Evolution of host resistance and damage-limiting mechanisms to an emerging bacterial pathogen

**DOI:** 10.1101/486365

**Authors:** Camille Bonneaud, Luc Tardy, Mathieu Giraudeau, Geoffrey E. Hill, Kevin J. McGraw, Alastair J. Wilson

## Abstract

Understanding how hosts minimise the cost of emerging infections has fundamental implications for epidemiological dynamics and the evolution of pathogen virulence. Despite this, few experimental studies conducted in natural populations have explicitly tested whether hosts evolve resistance, which prevents infections or reduces pathogen load through immune activation, or tolerance, which limits somatic damages without decreasing pathogen load. In addition, none have done so controlling for the virulence of the pathogen isolate used, despite critical effects on host responses to infection. Here, we conducted an experimental inoculation study to test whether eastern North American house finches (*Haemorrhous mexicanus*) have evolved resistance or tolerance to their emerging bacterial pathogen, *Mycoplasma gallisepticum,* using 55 distinct isolates of varying virulence. First, we show that peak pathogen loads, which occurred around 8 days post-inoculation, did not differ between experimentally inoculated finches from disease-exposed (eastern) *versus* unexposed (western) population. However, pathogen loads subsequently decreased faster and to a greater extent in finches from exposed populations, indicating that they were able to clear the infection through adaptive immune processes. Second, we found no between-population difference in the regression of clinical symptom severity on pathogen load; if tolerance had evolved then the slope of this regression is predicted to be shallower (less negative) in the exposed population. However, finches from exposed populations displayed lower symptom severity for a given pathogen load, suggesting that damage-limitation mechanisms have accompanied the evolution of immune clearance. These observations show that resistance and damage-limitation mechanisms - including, but not limited to the standard conceptualisation of tolerance - should not be seen as mutually exclusive. Nevertheless, we propose that host resistance is especially likely to evolve in response to pathogens such as *M. gallisepticum* that require virulence for successful infection and transmission.

## Introduction

Hosts can alleviate the costs of infection by evolving one of two distinct - though not necessarily mutually exclusive - strategies [1, 2]. They can evolve resistance, which serves to reduce the establishment of infectious pathogens and/or to clear pathogens following establishment [3, 4]. Alternatively, hosts can evolve tolerance, which serves to mitigate collateral somatic damage caused by the infection without reducing pathogen load [5–7]. Whether and when hosts evolve resistance or tolerance in response to emerging pathogens has far-reaching consequences for predicting virulence evolution and epidemiological dynamics [8], as well as for the design of novel pharmacological treatments [9, 10]. Despite this, few experiments have been performed to investigate the relative importance of resistance *versus* tolerance in evolved responses to naturally emerging pathogens.

Hypotheses based on the evolution of resistance *versus* tolerance make two contrasting predictions. First, if resistance (but not tolerance) has evolved, hosts will display reduced pathogen load during an infection relative to non-evolved hosts [8, 11, 12]. Second, if tolerance (but not resistance) has evolved, hosts will have a shallower regression of clinical symptom severity on pathogen load, since tolerant hosts are better able to mitigate the impact of an increasing pathogen load [12]. However, given these contrasting predictions, it is important to note that damage-limitation mechanisms could evolve in conjunction with resistance (e.g. by limiting immunity or initiating repair) [13, 14]. Thus, resistance and tolerance-mechanisms need not be mutually exclusive, and evidence for the evolution of one is not necessarily evidence against evolution of the other.

There have been few experimental tests of the predictions for the evolution of resistance and tolerance in response to naturally emerging pathogens, and the handful of studies to date have also yielded rather mixed conclusions. For example, strong evidence for the evolution of resistance comes from observations of the epidemic of myxoma virus in European rabbits (*Oryctolagus cuniculus*) in Australia [15, 16]. Following initially dramatic population declines, at 7 years post-outbreak rabbits from disease-exposed populations displayed mortality rates of only ∼25% in response to experimental infection. In contrast mortality rates of over 88% were found in unexposed wild and domestic rabbits [17, 18]. Subsequent work confirmed that reduced mortality was mediated by evolution of innate and cellular immune responses leading to significantly reduced pathogen loads [19]. By contrast, the endemic Hawaiian bird, Hawai‘i ‘Amakihi (*Chlorodrepanis virens*), was suggested to have evolved tolerance to *Plasmodium relictum* following the pathogen’s introduction to the archipelago in the 1930s [20]. Experimentally infected ‘Amakihi from high-altitude sites (where lower temperatures limit mosquito numbers and malaria parasite development; [20, 21]), displayed significantly higher mortality and weight loss than did individuals from low-altitude sites. Crucially, however, there was no significant difference in pathogen load [22].

Here, we examined the role of resistance *versus* tolerance in evolutionary responses of North American house finches (*Heamorhous mexicanus*) to the emerging, conjunctivitis-causing bacterium *Mycoplasma gallisepticum* following its jump from poultry in 1994 [23, 24]. Several previous experiments on this system have yielded apparently contradictory conclusions. Bonneaud et al. [25] concluded that resistance had evolved from standing genetic variation within 12 years of outbreak: finches from disease-exposed populations displayed reduced pathogen load following infection with a virulent, contemporary 2007 isolate. By contrast, Adelman et al. [26] concluded that tolerance had evolved: finches from disease-exposed populations in 2010 had similar pathogen load, but reduced symptoms (conjunctival swelling) relative to finches from unexposed populations and following inoculation with a low-virulence bacterial isolate collected at epidemic outbreak (i.e. in 1994). Although we reiterate that tolerance and resistance need not be mutually exclusive, further experiments are clearly needed to characterize their relative role in the response to this emerging pathogen. In addition, because support for tolerance evolution arises in part from a lack of significant differences in pathogen load (which are expected under resistance [7]), here we use a greater number of host individuals and of pathogen isolates of varying levels of virulence to reduce the possibility of type 2 error.

We conducted a large-scale infection experiment using 108 naïve house finches from disease-exposed (N=51) and unexposed (N=57) populations, and 55 bacterial isolates collected from the epidemic outbreak (1994) and during the subsequent 20 years (until 2015). After emergence near Washington D.C. in 1994, *M. gallisepticum* spread throughout the entire eastern US range of house finches within three years, killing millions of birds [23, 27]. While it later spread through much of the native western range (between 2000 and 2010; [28, 29]) some populations remain unexposed to date (e.g. in Arizona; [30]). We have shown previously that *M. gallisepticum* increased in virulence over the course of the epidemic, and that house finches from exposed populations displayed less severe symptoms than those from unexposed populations [31]. In this study, we test the key contrasting predictions set out above to determine whether this host evolutionary response is principally attributable to changes in resistance or tolerance.

First, if finches from exposed populations have evolved resistance, we would expect them to display lower pathogen loads during infection than birds from unexposed populations (i.e. populations that have not evolved resistance). Specifically, because resistance to *M. gallisepticum* is thought to be mediated through the ability to mount a cell-mediated immune response [32], given evolved resistance, finches from exposed populations are expected to show reduced pathogen load from approximately 2 weeks post-infection (i.e. the time required to mount a pathogen-specific immune response). By contrast, if tolerance alone has evolved, we predict no differences in pathogen load over the course of the month-long infection experiment, but predict that the relationship between symptom severity and pathogen load will be shallower in birds from exposed populations [12].

## Results

Over the course of the experiment, the median peak bacterial load observed across the 108 individual birds was 78 bacteria per host cell. There was marked variation around this median (IQR of 42-154, total range of 1-522 bacteria.cell^−1^), arising in part from effects of bacterial isolate identity. Specifically, a mixed model analysis revealed that isolate identity explained 19% of the variance in peak load. Notably, isolates sampled from early in the epidemic (1994-2003) generated similar peak loads in birds from exposed and unexposed populations, whereas differences in bacterial load between populations tended to be of greater magnitude with infection by later sampled isolates (2007-2015) (Figure 1). Overall, however, there was no directional pattern and average peak load was similar in birds from exposed and unexposed populations (mixed GLM; population effect (unexposed relative to exposed) ± se = −0.10 ± 0.29, *χ*^2^ = 0.11, df = 1, p =0.74).

**Figure 1.**
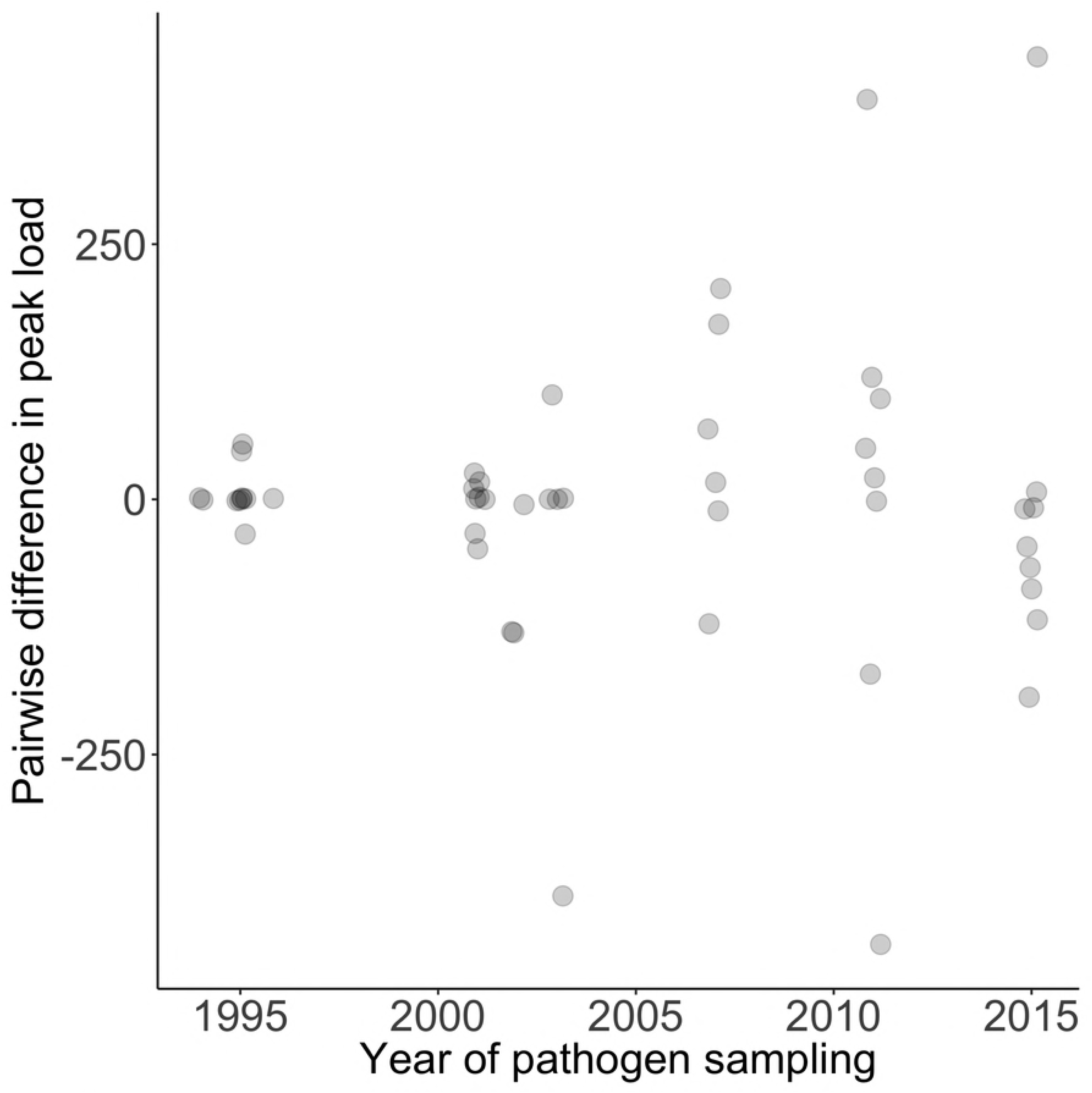
Population differences in peak pathogen load. We show differences in peak pathogen load (measured in number of bacteria per host cells) at the site of infection between pairs of birds from unexposed *versus* exposed pathogen load that were inoculated with the same bacterial isolate (N = 51 pairs), as a function of the year of isolate sampling.

The absence of a population difference in peak load is not necessarily incompatible with evolved resistance if pathogen loads are peaking prior to the time when genetically resistant birds are able to mount an effective immune response. Here bacterial loads were highest (on average) in birds from both populations at 8 days post-inoculation (dpi) and thereafter declined significantly (mixed GLM; main dpi effect: estimate ± se = −0.07 ± 0.009, *χ*^2^ = 54.7, df = 1, p < 0.0001). However, there was a significant population by dpi interaction (estimate ± se = 0.07 ± 0.02, *χ*^2^ = 13.9, df = 1, p < 0.0002), with birds from exposed populations clearing the pathogen approximately 3 times faster than those from unexposed populations (Figure 2). Differential clearing rates are such that birds from exposed populations have a 4-fold lower bacterial load than birds from unexposed populations by 28 dpi. This finding is consistent with the hypothesis that genetic resistance through acquired immunity has evolved in exposed populations.

**Figure 2.**
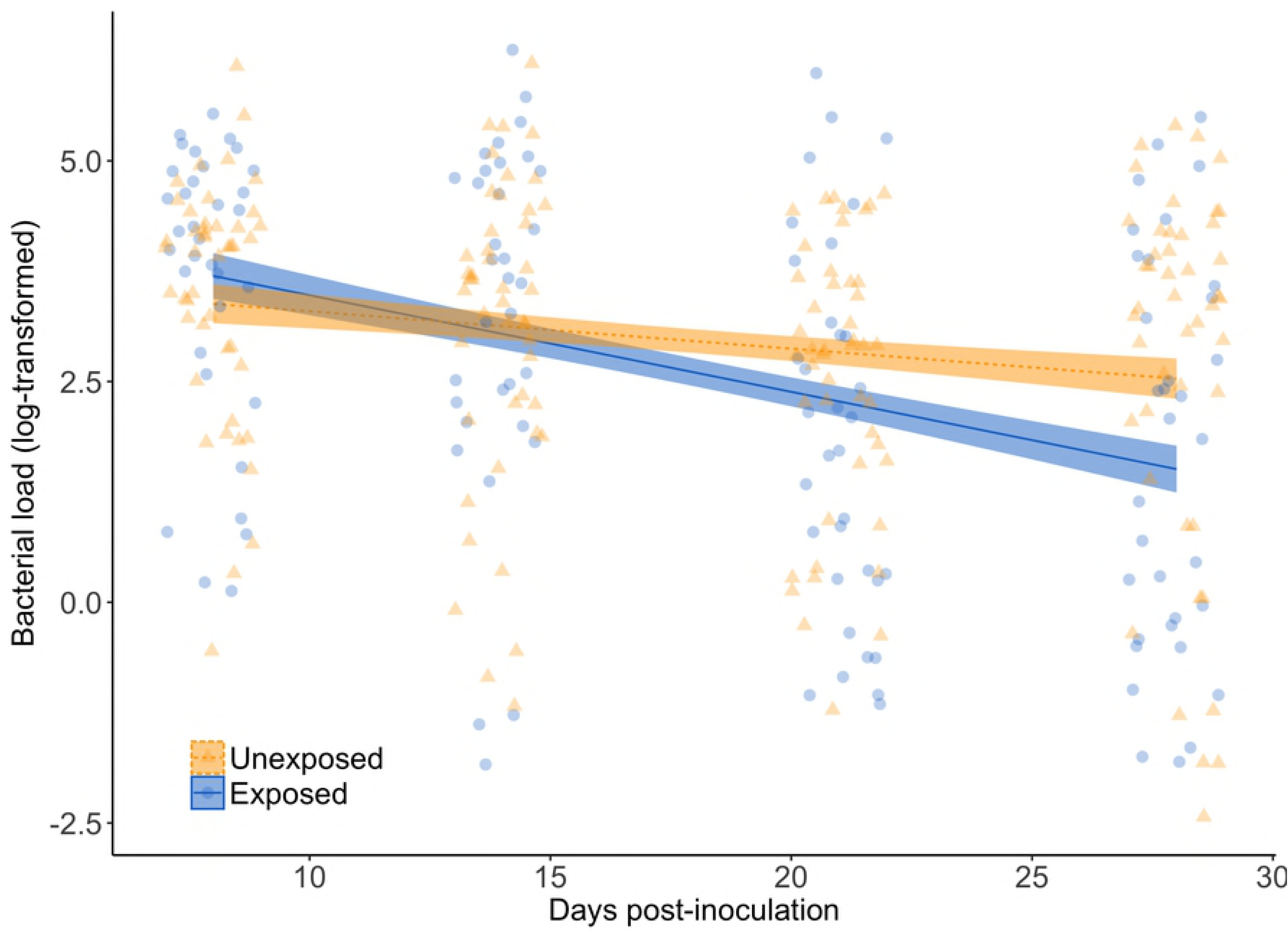
Changes in pathogen load over the experiment. We show pathogen load (log-transformed) at 8, 14, 21 and 28 dpi for birds from exposed (blue) and unexposed (orange) populations. Raw values are shown as triangles (exposed) or circles (unexposed populations); lines are predicted from the model (solid = exposed; dashed = unexposed), with standard errors represented by ribbons.

The key symptom of *M. gallisepticum* infection in house finches is conjunctivitis, which, when severe, causes blindness and death in the wild through starvation or predation [33–35]. Using the area of conjunctival swelling to measure clinical symptom severity, we found no clear support for the hypothesis that tolerance has evolved. Across all observations over the course of the experiment, the average measure of peak conjunctival swelling was 64 34 mm2. To account for changes in pathogen load over time, we calculated the integral of pathogen load over the course of the experiment. Peak swelling increased with the integral of pathogen load (mixed GLM; pathogen load main effect: estimate ± se = 0.25 ± 0.13, *χ*^2^ = 24.4, df = 1, p < 0.0001), but the slope of this regression did not differ between birds from exposed *vs.* unexposed populations (population × integral of pathogen load interaction effect: estimate ± se = −0.03 ± 0.08, *χ*^2^ = 0.4, df = 1, p = 0.73), as predicted under the hypothesis that tolerance alone had evolved (Figure 3). Nonetheless, birds from exposed populations have 10% lower clinical symptoms for a given pathogen load (population main effect: estimate ± se = 0.21 ± 0.04, *χ*^2^ = 6.3, df = 1, p = 0.012). Thus, while there is no evidence that (slope) tolerance differs between populations, our results do suggest that some mechanism(s) to limit immune damage have evolved in tandem with resistance. Note that results were qualitatively equivalent when using peak pathogen load rather than integral of pathogen load (see supplementary information).

**Figure 3.**
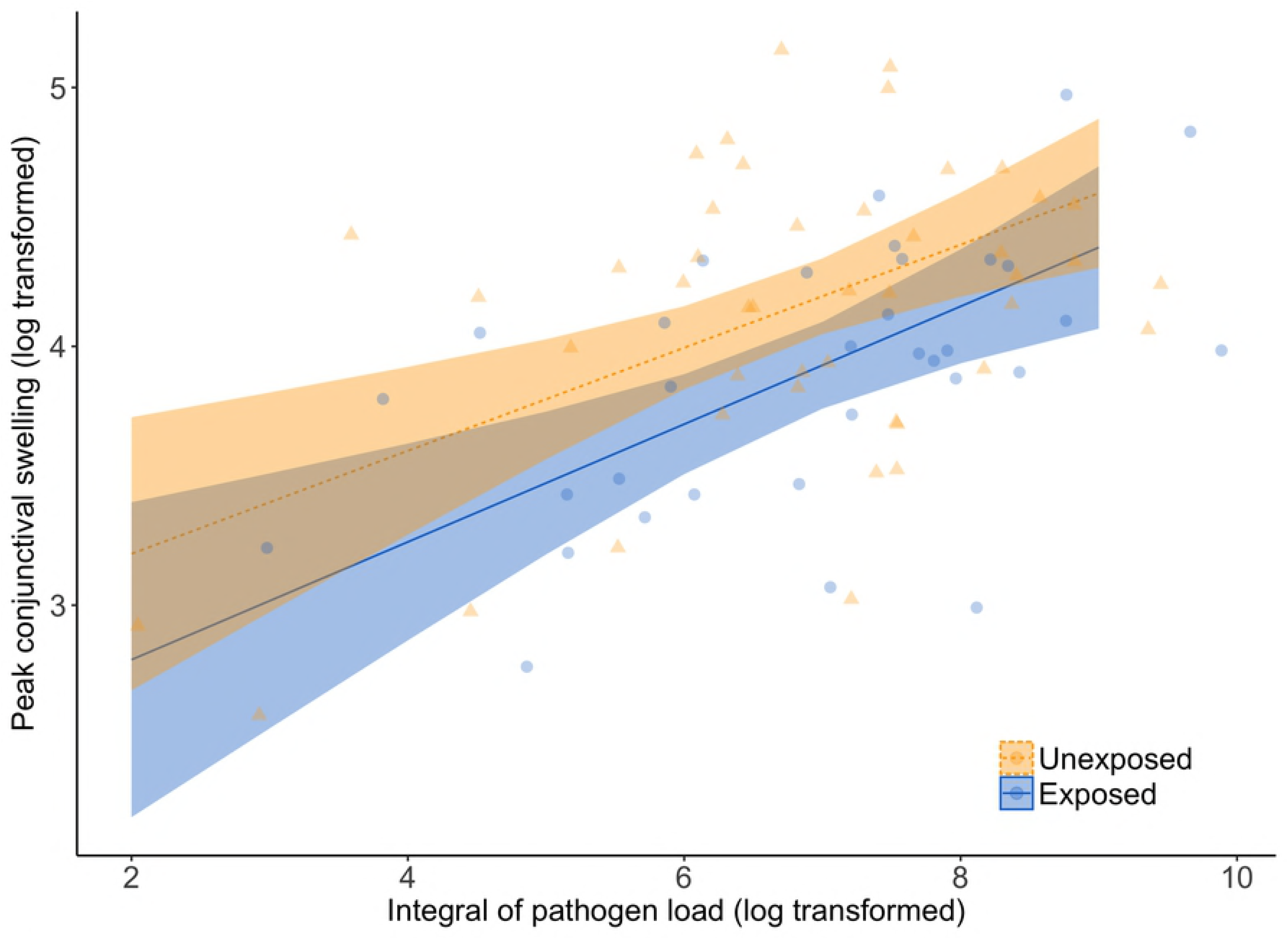
Association between pathogen load and clinical symptom severity. We show peak conjunctival swelling (log transformed) as a function of the integral of pathogen load over ythe course of the experiment(log-transformed) for birds from exposed (blue) and unexposed (orange) populations. Raw values are shown as triangles (exposed) or circles (unexposed populations); lines are predicted from the model (solid = exposed; dashed = unexposed), with confidence intervals represented by ribbons.

## Discussion

To test whether house finch hosts from disease-exposed populations have evolved resistance or tolerance to infection to the emerging bacterial pathogen *M. gallisepticum,* we conducted an inoculation experiment of house finches from disease-unexposed and exposed populations using 55 isolates collected over a 20-year period from epidemic outbreak. We found that birds from exposed and unexposed populations had comparable peak pathogen loads, which were maximal at 8 days post-inoculation in both cases. However, thereafter birds from previously exposed populations cleared the pathogen more rapidly and to a greater extent during our experiment. We interpret these patterns as clear evidence that, in the exposed populations, hosts have evolved resistance in response to the emerging pathogen *M. gallisepticum*.

In contrast to the evidence supporting the hypothesis of evolved resistance, evidence for the evolution of tolerance in the exposed finch population was equivocal. Notably, the gradient of the regression of symptom severity on pathogen load was comparable in birds from exposed and unexposed populations. Because we did not observe the predicted differences in the response slopes of exposed *vs*. unexposed host populations, our results are not consistent with the hypothesis that tolerance to *M. gallisepticum* has evolved. Nonetheless, birds from exposed populations did exhibit lower clinical symptoms across the range of pathogen loads experienced (i.e. the intercept of the regression line is lower than in unexposed populations). This pattern suggests that some mechanism(s) to limit damage (i.e. symptom severity) has evolved, although whether this should be interpreted as tolerance is perhaps a matter of perspective. The ambiguity arises because, while we have adopted a standard ‘range tolerance’ concept (*sensu* [36]), ‘point tolerance’ approaches in which, for example, tolerance differences are inferred from correlates of host fitness differences at a single pathogen load, are also widely used in the literature (see e.g., [37] and references therein for further discussion).

In this study, we found a difference in reaction norm intercept, but not slope, between unexposed and exposed finch populations. Strictly, this provides evidence for the evolution of ‘point’, but not ‘range’ tolerance. Whether this semantic distinction adds much biological insight is arguable, but what does seem clear is that any change in tolerance that has occurred has been accompanied by a change in resistance. Our finding that bacterial loads only started to differ between exposed and unexposed finch populations after 14 dpi, as predicted if clearance occurs through cell-mediated immunity [32], may also help to reconcile the somewhat divergent results of previous studies in house finches [25, 26]. For instance, Adelman et al. [26] concluded an important role for tolerance, but not resistance, because birds inoculated with *M. gallisepticum* displayed lower peak eye lesion scores in finches from exposed populations, despite no significant difference in bacterial load. However, load was measured at peak, which occurred before 10 dpi (as per this study) and at which point we would not expect a detectable increase in the protective immune activity in birds from exposed populations. Nevertheless, Adelman et al. [26] did not find evidence for a subsequent between-population differences in pathogen clearance, an effect which may be attributable to a restricted sample size typical of previous studies (see also [25]). Finally, we now know that there is substantial among-isolate variation in virulence [31], and that differences in symptom severity between exposed and unexposed host populations are more apparent under infection with late-epidemic bacterial isolates. It is therefore possible that use of a single early-epidemic isolate may have contributed to the previous conclusion that resistance has not evolved [26]; indeed this would be expected if the low virulence of the isolate used [31] barely elicited a protective immune response in birds from exposed populations.

Taken together, the results of our study highlight that future tests of resistance *versus* tolerance evolution in response to naturally emerging pathogens require: (i) analyses of pathogen load over a sufficient infection duration to encompass the consequences of both innate and adaptive immune processes; (ii) inoculations with sufficient numbers of pathogen isolates taken from varying time-points of the host-pathogen interaction; (iii) a clear operational definition of tolerance; and, (iv) a greater recognition that while resistance and tolerance can be viewed as distinct host defence strategies, this does not mean they must be either mechanistically independent or mutually exclusive. Indeed, immune cells are increasingly recognized as playing a dual role in resistance and damage-limitation processes in the broad sense [38, 39]. We do, however, suggest that divergent concepts of tolerance in the literature necessitate some caution when interpreting results. If *range tolerance* has evolved, the prediction is that the regression of pathogen load on symptom severity will be reduced, not just that we should find lower symptom severity for a given pathogen load overall. This prediction is not realised in this case, but this certainly does not mean that there is no value in future studies addressing the role of damage-limitation mechanisms that curtail and resolve immune responses to prevent autoimmunity, remove cellular debris and stimulate tissue repair and regeneration [39].

More generally, it has been suggested recently that the evolution of damage-limitation mechanisms is necessarily indicative of an important role of tolerance in host evolution. We view this as potentially problematic for two reasons. First, damage-limitation can clearly evolve in tandem with immune clearance in which case it can be viewed, at least in part, as a consequence of host resistance. Second, not clearly distinguishing between resistance and tolerance, whether as a consequence of an inadequate study design or because the two concepts are not actually biologically distinct in a given system, will negatively impact the ability to predict co-evolutionary dynamics. For instance, although resistance is implicated in antagonistic host-pathogen coevolution [40, 41], it is often noted that the emergence of tolerance should benefit both parties, allowing interactions to evolve towards commensalism [8, 42]. In fact, this latter prediction is likely contingent on the assumption that virulence is a by-product of pathogen replication rates [8, 41, 43–46]. We argue that when virulence is required for pathogen success [47], the evolution of host tolerance will actually select for increased pathogen virulence.

Crucially in the current context, *M. gallisepticum* requires virulence to establish infection and transmit between hosts. This is because transmission occurs through ocular fluid exudates [48] and so depends on the bacterium causing a misdirected inflammatory response that disrupts the mucosal surface of the conjunctiva and respiratory tract [49–52]. Given that high-virulence is broadly expected to favour the evolution of host resistance, while low-virulence should favour tolerance [53], it is intuitive that obligately virulent pathogens should lead to the evolution of resistance. In finches, resistance to *M. gallisepticum* via an effective cell-mediated immune response does seem to have evolved, but has likely been accompanied by the ability to resist the pathogen-driven activation of an inflammatory response [26, 54]. This interpretation is consistent with our finding that the slopes of the relationships between pathogen load and clinical symptoms severity (i.e. ‘range’ tolerance) were equivalent between finches from disease-exposed and unexposed populations, but those from the latter displayed higher symptoms overall.

In conclusion, we provide evidence that house finches have evolved resistance following the infectious outbreak of the bacterial pathogen, *M. gallisepticum*, with finches from disease-exposed populations likely reducing pathogen load through acquired immune processes [25]. Further, however, we also found evidence to suggest that the ability to limit damage by the pathogen has evolved in tandem with resistance. Although tolerance mechanisms have been typically defined and conceptualised as an evolutionary alternative to resistance [6], it is not obvious why resistance and tolerance need be antagonistic. Indeed, under ‘classic’ tolerance as well as when tolerance mechanisms accompany the evolution of resistance, the key effect is to reduce the damage caused by the infection. Thus, given that tolerance decreases the costs of infection irrespective of whether or not it evolves with resistance, we suggest that a more general view of tolerance might be sensible. Under this more general framework, the evolution of tolerance and resistance can be considered as two complementary strategies, which can evolve together to fight infection and reduce damage.

## Material and Methods

### Capture and housing

Wild hatch-year house finches from populations that have never been exposed to *M. gallisepticum* (unexposed populations) were captured in urban areas and in suburban parks [55, 56] in Arizona over a two-week period of the summer 2015. We trapped, weighed and banded each bird with a numbered metal tag for individual identification (N=171; 93 males and 78 females). They were then immediately transported by car to aviaries at Arizona State University, where they were housed for the remainder of the experiment. On arrival, we sampled blood from each bird by brachial venipuncture (60 μL of whole blood) and also took a choanal swab. A lack of prior infection was confirmed for each bird by screening blood plasma samples for anti-*M. gallisepticum* antibodies using a serum plate agglutination assay [57]. Absence of current infection was verified using the choanal swabs in PCR amplification of *M. gallisepticum* DNA [58]. No prior or current infections were detected (as expected given no documented reports of *M. gallisepticum* from this area of Arizona [30]).

During the same time period, we also caught hatch-year house finches from populations known to have been exposed to *M. gallisepticum* since the disease outbreak (exposed populations). These were captured from urban areas and suburban parks in Alabama (see [31]). Birds were similarly banded weighed, and sampled for blood and choanal swabs. They were then immediately transported by car to aviaries at Auburn University, where they were housed separately in the same conditions as in the aviaries in Arizona and tested for prior and current infection as described above. Birds positive for either test were released immediately, while the remaining individuals (N=53; 24 males and 29 females) underwent a 30-day quarantine period, before being transported by car to the aviary at Arizona State University.

Following arrival at Arizona State University, 108 birds (51 birds from the exposed populations and 57 birds from the unexposed populations) were haphazardly selected for us in the present study (the remaining 116 individuals being used in another experiment). They were then allowed to acclimate in the aviaries, with *ad libitum* food and water, for one month prior to experimental inoculation. During this period no birds displayed any sign of infection with other diseases but all were treated prophylactically for *Trichomonas gallinae* and *Isospora spp* in the first few weeks of captivity as a precaution.

### Experimental inoculation

We inoculated each of the birds with one of fifty-five *M. galliseptum* isolates sampled over the course of the epidemic (N total inoculated =108, consisting of 51 birds from exposed populations inoculated with 51 isolates and 57 birds from unexposed populations inoculated with 55 isolates; 2 isolates were each inoculated into 2 birds from unexposed populations). Isolates were administered via 20 μL of culture containing 1 × 10^4^ to 1 × 10^6^ colour changing units/mL of *M. galliseptum* in both eyes. We then photographed the right and left eyes at 0, 6, 13 and 25 days post-inoculation (dpi) to monitor the development of conjunctival swelling. This was done by measuring the area of the conjunctiva swelling as the area of the outer ring minus the area of the inner ring (see [30]). Measurements were blind with respect to the isolate inoculated and the population of origin of the bird. Mean conjunctival swelling was then determined for a bird by averaging means of both eyes across the four dpi specific measurements. The experiment was stopped at 35 dpi and all birds were euthanized as stipulated by Home Office (UK) licencing. Protocols were approved by Institutional Animal Care and Use Committees (IACUC) of Auburn University (protocol # PRN 2015-2721) and of Arizona State University (protocol #15-1438R), and by Institutional Biological Use Authorizations to Auburn University (# BUA 500).

### Pathogen load

Bacterial load was measured by quantitative amplification of *M. gallisepticum* DNA from conjunctival and tracheal swabs obtained at 8, 14, 21 and 28 dpi. DNA was extracted using a QIAGEN DNeasy^®^ Blood and Tissue Kit according to the manufacturer’s standard protocol. For each sample, we ran a multiplex quantitative PCR of the *M. gallisepticum*-specific gene *mgc2*, which encodes a cytadhesin protein, and the house finch recombination-activation gene *rag1*, using an Applied Biosystems™ StepOnePlus™ Real-Time PCR system (see supplementary information). Each reaction contained: 2µl of sample genomic DNA template, 1 µl each of 10 µM mgc110-F/R and rag1-102-F/R primers (total 4ul), 0.5 µl each of 10 µM Mgc110-JOE and Rag1-102-6FAM fluorescent hydrolysis probes (total 1 µl), 10 µl of 2X qPCRBIO Probe Mix HI-ROX (PCR BIOSYSTEMS) and 3 µl Nuclease-free water (Ambion^®^). Reactions were then run at 95°C for 3 min, followed by 45 cycles of 95°C for 1s and 60°C for 20s. Samples were run in duplicate alongside serial dilutions of plasmid-based standards (range of standards for *mgc2*: 1.6×108 – 1.6×103 copies; range of standards for *rag1:* 8.0×107 – 8.0×102 copies). Amplification data was exported to LinRegPCR v.2017.1 for calculation of individual reaction efficiencies and quantification of low-amplification samples [59, 60]; between run variation was normalised using Factor qPCR v.2016.0 [61], with plasmid standard serial dilutions used for factor correction.

### Statistical analyses

All statistical analyses were conducted in R 3.3.2 [62] using linear mixed effect models fitted in lme4 [63], and figures were made using ggplot2 [64]. Response variables (pathogen load, conjunctival swelling data) were natural log-transformed to better meet the assumptions of residual normality. To determine whether peak pathogen load differed between disease-exposed and unexposed host populations, we ran a mixed effects model with ln(pathogen load) as the response term, host population (exposed vs unexposed), dpi (as a continuous covariate) and their interaction as explanatory terms. Bacterial isolate identity and bird identity were both included as random effects. Finally, we tested for population differences in the association between pathogen load and clinical symptoms severity using a second mixed effects model. Here ln(peak conjunctival swelling) was the response term, with host population, integral measure pathogen load over the course of the experiment and their interactions as fixed effects. Bacterial isolate identity was again included as a random term but not individual identity (since each bird was only represented once). We note that log transformations stabilize variance and in this case that risks removing (or at least reducing) any signature of a population × pathogen interaction on symptom severity. Since this interaction is key to our hypothesis testing (i.e. we predict a steeper regression of symptom severity on pathogen load in unexposed populations if tolerance has evolved) we re-ran this second analysis using untransformed conjunctival swelling data. Since conclusions were unaltered we elected not present that analysis here (but see results in supporting information).

## Acknowledgments

This research was supported by a Natural Environment Research Council standard grant to C.B. and A.W. (NE/M00256X). We thank A. Russell for helpful discussion and constructive comments on the manuscript. We are grateful to James S. Adelman and Dana M. Hawley for helpful discussions. We thank M. Staley for growing and shipping the pathogen isolates, M. Cooke for assisting with bird captures in Arizona, A. Santos, W. R. Hood and the undergraduates in the Hood lab for assisting with bird captures in Alabama, and A. K. Ziegler for assisting with the experiment in Arizona.

## Author Contributions

CB conceived and designed the study. GEH, KJM, MG, CB obtained the animals and/or bacterial isolates. MG, KJM, LT conducted the experiment. LT conducted the molecular work. CB and AW analysed the data and wrote the paper.

## Data Availability Statement

All data will be made available on Dryad Digital Repository (https://datadryad.org).

